# LightRoseTTA: High-efficient and Accurate Protein Structure Prediction Using an Ultra-Lightweight Deep Graph Model

**DOI:** 10.1101/2023.11.20.566676

**Authors:** Xudong Wang, Tong Zhang, Guangbu Liu, Zhen Cui, Zhiyong Zeng, Cheng Long, Wenming Zheng, Jian Yang

## Abstract

Accurately predicting protein structure, from amino acid sequences to three-dimensional structures, is of great significance in biological research. To tackle this issue, a representative deep big model, RoseTTAFold, has been proposed with promising success. Here, we report *an ultra-lightweight deep graph network*, named *LightRoseTTA*, to achieve accurate and high-efficient prediction for proteins. Notably, three highlights are possessed by our LightRoseTTA: **(i) high-accurate** structure prediction for proteins, being *competitive with RoseTTAFold* on multiple popular datasets including CASP14 and CAMEO; **(ii) high-e**ffi**cient** training and inference with an ultra-lightweight model, costing *only one week on one single general NVIDIA 3090 GPU for model-training* (vs 30 days on 8 high-speed NVIDIA V100 GPUs for RoseTTAFold) and containing *only 1*.*4M parameters* (vs 130M in RoseTTAFold); **(iii) low dependency** on multi-sequence alignments (MSA, widely-used homologous information), achieving *the best performance on three MSA-insu*ffi*cient datasets: Orphan, De novo, and Orphan25*. Besides, our LightRoseTTA is *transferable* from general proteins to antibody data, as verified in our experiments. We visualize some case studies to demonstrate the high-quality prediction, and provide some insights on how the structure predictions facilitate the understanding of biological functions. We further make a discussion on the time and resource costs of LightRoseTTA and RoseTTAFold, and demonstrate the feasibility of lightweight models for protein structure prediction, which may be crucial in the resource-limited research for universities and academy institutions. *We release our code and model to speed biological research*.

## 1. Introduction

Proteins play significant roles in the biological activities of living organisms, including transporting substances, participating in immunity, and regulating hormones. As the function of proteins is largely determined by the structure, making clear three-dimensional structures would give important insights into understanding biological mechanisms in cellular processes and facilitate interventions (e.g. drug development) through protein function modulation. The previous conventional methods of experimental structure determination, using either nuclear magnetic resonance (Thompson et al., 2020), X-ray crystallography (Jaskolski et al., 2014), or cryo-electron microscopy (Bai et al., 2015), can be rather time-consuming and expensive. Hence, it has become a hot research topic which is to leverage machine learning algorithms to predict three-dimensional protein tertiary structures from amino acid sequences.

Tremendous efforts have been made for protein structure prediction in the past decade. In the early stage, one main technical line primarily considers physical chemistry principles of proteins in structure prediction, e.g. assuming the lowest-energy conformation for folded proteins. However, it suffers from the difficulties of determining a widely applicable energy function as well as the exhausting search in the huge conformational space. As a result, limited prediction performances were achieved. In recent years, with the ever-growing sequences in protein databases such as UniProt Knowledge base (UniProtKB) (Suzek et al., 2015; Mirdita et al., 2017), deep learning algorithms have been introduced for protein structure prediction (Senior et al., 2020; Roy et al., 2010; Shindyalov et al., 1994; Yang et al., 2020; Hiranuma et al., 2021; Marks et al., 2012). Generally, the main-stream deep models utilize information from both template structures and homologous sequences, and further design neural networks with rather deep architecture to learn the co-evolved features. Following this pipeline, RoseTTAFold (Baek et al., 2021), as one of the representative models, achieves promising performance on the CASP14 dataset. However, the deep models heavily depend on multi-sequence alignment (MSA), and thus obtain poor prediction accuracy on those proteins with few or limited homologous sequences. To solve this issue, some MSA-free deep learning methods (Wang et al., 2022; Chowdhury et al., 2022), named trRosettaX-single and RGN2, have been proposed to predict structures by using only textual context of amino acid sequences without MSA, effectively improving the performance on the proteins with few homologous sequences.

Although significant success has been made by previous works, two critical issues still remain in promoting research and application of protein structure prediction. The deep big models suffer from high computational cost and resource burden. For instance, RoseTTAFold needs 30 days to be trained on 8 NVIDIA V100 GPUs. This limits the updating efficiency of the research and model development, meanwhile, deters most universities and academy institutions from participating in this research field. Second, a better MSA-dependency reduction strategy is urgently required to jointly perform well on both sufficient and insufficient homologous sequences. Although the existing MSA-free deep learning methods improve the prediction performance on proteins with insufficient homologous sequences, their results on other homology-sufficient proteins are not satisfying, finally leading to relatively low overall performance.

In this work, we describe a novel ultra-lightweight deep framework, named LightRoseTTA to accurately and efficiently predict 3D all-atom coordinates of proteins. Primarily, compared with RoseTTAFold, three outstanding advantages are possessed by our LightRoseTTA, (i) more competitive performance on protein structure prediction; (ii) the much smaller model size with only 1.4M parameters, costing much less computational time and resource in the model-training stage; (iii) the lower dependency on homologous sequences. For the model design, a backbone-to-all-atom architecture is proposed to accurately predict 3D all-atom coordinates by leveraging high backbone-sidechain dependency. Specifically, for the backbone conformation, a two-branch network, taking MSA, template, and atom-level information as inputs, is designed to model both inter-residue relationships and sidechain influences. Most critically, to compress the model while achieving high performance, we observe and leverage a constraint prior named backbone potential energy (BPE), normalizing bond lengths, bond angles and dihedrals, to guide the backbone generation. The BPE constraint effectively regularizes the parameter optimization space for backbone generation, enabling a significantly shallower architecture network to handle the complicated backbone conformation as validated in experiments. Moreover, to well handle proteins with both sufficient and insufficient homologous sequences, a new MSA dependency-reducing strategy is further designed during the model training.

We report the performance on the popular CASP14 dataset (critical assessment of techniques for protein structure prediction (Kryshtafovych et al., 2021)) and CAMEO dataset (continuous automated model evaluation (Robin et al., 2021)), as well as the homology-insufficient Orphan (Chowdhury et al., 2022), De novo (Chowdhury et al., 2022), Orphan25 (Wang et al., 2022) and Design55 (Wang et al., 2022) datasets. The experimental results demonstrate the more competitive performances (TM-score (Zhang and Skolnick, 2004) and GDT TS (Zemla, 2003)) of our ultra-lightweight LightRoseTTA compared with RoseTTAFold on CASP14 and CAMEO, and homolog-insufficient datasets Orphan, De novo and Orphan25. Furthermore, to check the feasibility of LightRoseTTA across different protein categories, e.g., general protein versus antibody, we transfer our LightRoseTTA to predict the antibody structure on Rosetta Antibody Benchmark (Ruffolo et al.). LightRoseTTA achieves promising performance on the most critical region–the third complementary determining region ring of the heavy chain (CDR-H3) for antibodies, verifying its effectiveness and good transferability for proteins of different categories. Moreover, we discuss the time and resource costs of LightRoseTTA and RoseTTAFold for protein structure prediction. Our LightRoseTTA has only 1.4M parameters, and can be well trained on a single GPU (NVIDIA RTX 3090) in only one week. In contrast, RoseTTAFold costs 30 days using 8 NVIDIA V100 GPUs. Our LightRoseTTA achieves the best performances in the most cases while costing far less time and resources for model learning.

## 2. Result

### Approach Summary

#### The whole framework

The whole framework of the proposed LightRoseTTA is illustrated in Fig. 1. Overall, it employs the backbone-to-all-atom architecture for 3D structure prediction, which first predicts the backbone and then obtains the all-atom structure by leveraging high backbone-sidechain dependency. For one given amino acid sequence, three aspects of information, i.e., MSAs (Waterhouse et al., 2018; Zhang et al., 2020), structure templates, and atom-level graphs, are first generated as inputs of the LightRoseTTA. Next, a two-branch network, consisting of the residue-level and atom-level branches, is constructed to model both inter-residue interaction and sidechain influence for backbone conformation. The residue-level branch first applies the co-evolution learning module to alternately update MSA and template information in a co-evolutional way. It outputs both residue features and paired inter-residue relation. Then, a hybrid convolution neural network (CNN) (Krizhevsky et al., 2012), performing convolutions by using both classic (unsymmetric) and symmetric convolutional kernels, is designed to learn pairwise inter-residue representation. We further treat residue features and inter-residue representation as nodes and edges, respectively, to construct a residue-level graph. Subsequently, a graph transformer (Shi et al., 2021) is employed to learn geometric representation by aggregating residue features according to inter-residue edges. In contrast to RoseTTAFold (Baek et al., 2021), this residue-level branch has two notable differences: (1) possessing a rather shallow architecture of much fewer coevolution blocks in the co-evolution learning module; (2) designing the symmetric convolutional kernels in the hybrid CNN to preserve the intrinsic symmetry of pairwise attributes (e.g., inter-residue dihedrals *ω*: C_*α*_-C_*β*_-C_*β*_-C_*α*_).

**Figure 1.**
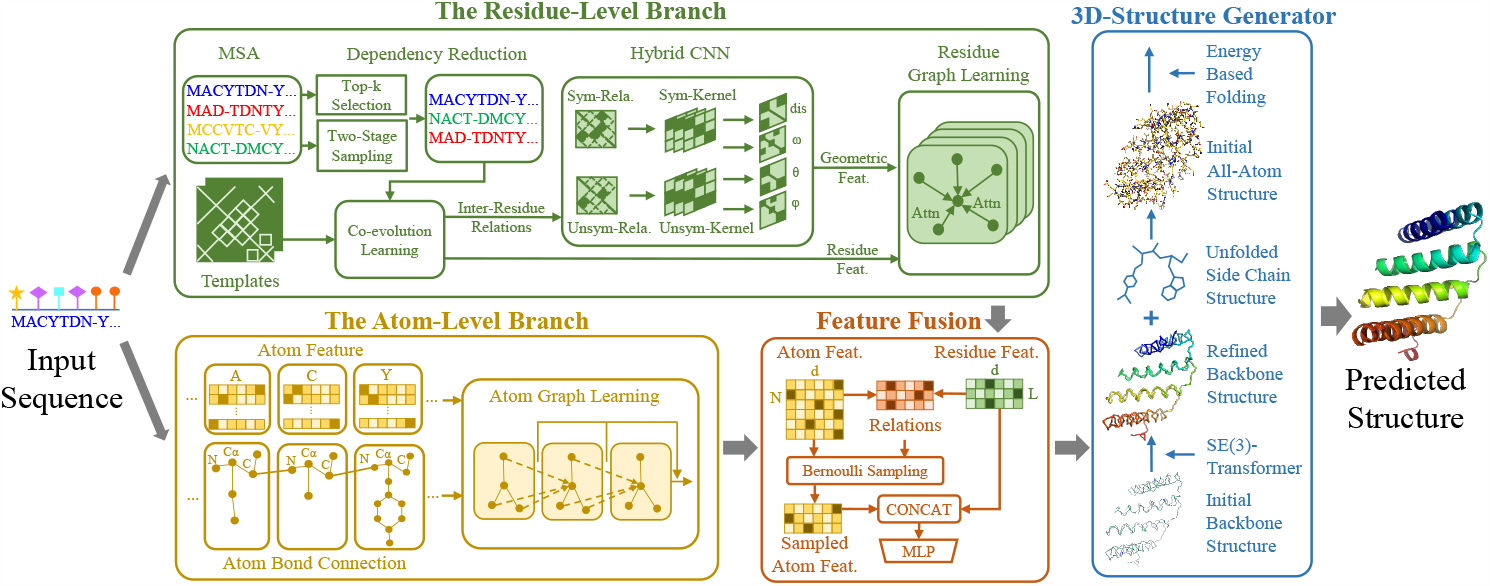
The architecture of LightRoseTTA for protein structure prediction. LightRoseTTA consists of two branches, the residue-level branch and the atom-level one. For one given amino acid sequence, three aspects of information, i.e., MSA features, structure templates, and atom-level graphs, are first generated as the inputs. The residue-level branch uses three hierarchical modules, i.e., the co-evolution learning module, the hybrid CNN module, and the residue graph learning module, to learn residue-level features. The other atom-level branch employs the atom graph learning module to aggregate features of atoms in both backbone and sidechains, resulting in atom-level representation. Next, the residue-level and atom-level features are fused through variational learning to refine the backbone information. Finally, the fused features are sent into the 3D-Structure generator to predict the 3D all-atom structure.

The other atom-level branch additionally takes the sidechain effects into account, and constructs the atom-level graph by regarding atoms and bonds as nodes and edges. The atom-level representation is then learned through graph neural network (GNN) (Morris et al., 2019) by aggregating features of atoms in both backbone and sidechains. The atom-level branch is fused into the residue-level branch through variational learning to inject sidechain awareness into backbone generation. Hence, the residue-level representation can be further refined by adaptively considering the influence of atom-level variation. After the two-branch fusion, the resulting features are fed into the 3D-Structure generator for both backbone and all-atom structure prediction. Concretely, the coordinates of the backbone atoms (N, C_*α*_, C) are first predicted through the SE(3)-Transformer (Fuchs et al., 2020). To further obtain the all-atom structure, next, the sidechain atoms are bonded to the backbone and optimized based on the all-atom potential energy.

For the model training, a constraint prior named backbone potential energy (BPE) is specifically proposed for backbone prediction. The BPE effectively regularizes the parameter optimization space, and enables a pretty shallow model to predict accurate backbone structures. Moreover, to make LightRoseTTA robust against homologous sequences, a new MSA dependency-reducing strategy is further designed during the model training. In the training stage, the whole architecture can be optimized in an end-to-end manner through gradient backpropagation.

#### Predicting protein structures in CASP14 and CAMEO

To evaluate the performance of the LightRoseTTA for protein structure prediction, we first compare it with those methods named RoseTTAFold (Baek et al., 2021) and ProFold (Ju et al., 2021) on the popular CASP14 (Kryshtafovych et al., 2021) and CAMEO (Robin et al., 2021) datasets. For performance measurement, the widely adopted TM-score (Zhang and Skolnick, 2004) and GDT TS (Zemla, 2003) (the higher the better for both of them) are used as the metrics for protein structure prediction. The experimental results are shown in Fig. 2a and Fig. 2b. Overall, our LightRoseTTA achieves competitive prediction performances on both CASP14 and CAMEO datasets. For the metric of TM-score, it outperforms RoseTTAFold by obtaining 0.004 (0.755 of LightRoseTTA vs 0.751 of RoseTTAFold) performance gain on CASP14 and achieving 0.002 (0.772 of LightRoseTTA vs 0.77 of RoseTTAFold) higher performance on CAMEO. Meanwhile, LightRoseTTA outperforms ProFold on both CASP14 and CAMEO with the performance improvements of 0.033 and 0.025, respectively. For the other metric of GDT TS, similar with the comparison results of TM-score, LightRoseTTA outperforms RoseTTAFold on CASP14 (70.34 of LightRoseTTA vs 70.07 of RoseTTAFold), and on CAMEO (68.78 of LightRoseTTA vs 68.3 of RoseTTAFold). Also, LightRoseTTA performs better than ProFold with the GDT TS improvements of about 2.78 and 2.69 on CASP14 and CAMEO, respectively. Moreover, we visualize several examples of LightRoseTTA-predicted structures in Fig. 2c-e, which intuitively show the high structural consistency between the predicted results (red) of our LightRoseTTA and the structure obtained through experiments (gray). Furthermore, we compare the prediction results of LightRoseTTA and RoseTTAFold with respect to the proportion of secondary structures (helices, beta-fragments and loops), and the protein length of CAMEO proteins in Supplementary Section 4, with the results shown in Supplementary Fig. 2.

**Figure 2.**
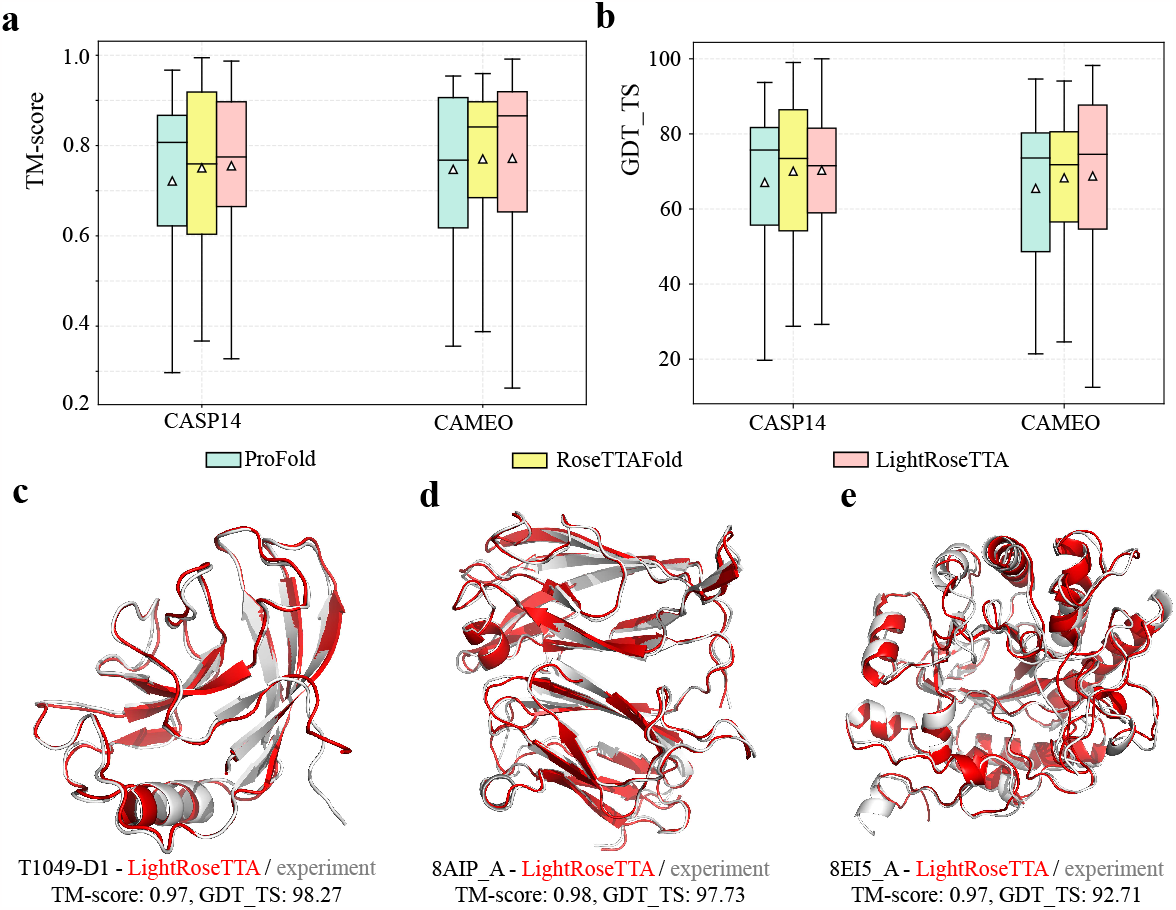
The performance comparison on the CASP14 and CAMEO datasets and the visualization examples of LightRoseTTA’s prediction. **a**, The performance (TM-score) on the CASP14 and CAMEO datasets. For each box in the figure, the center line, bottom line, and top line represent the median, first quartile, and third quartile, respectively. The horizontal lines along the top and bottom edges represent the maximum and minimum observations. Besides, the white triangle represents the average value. On the CASP14 dataset, the mean value of TM-score for ProFold, RoseTTAFold, and LightRoseTTA are 0.722, 0.751, and 0.755, respectively. For CAMEO, the value of three methods are 0.747, 0.77, and 0.772, respectively. **b**, The performance (GDT TS) on the CASP14 and CAMEO datasets. For CASP14, the mean value of GDT TS for ProFold, RoseTTAFold, and LightRoseTTA are 67.56, 70.07, and 70.34, respectively. For CAMEO, the value of three methods are 66.09, 68.3, and 68.78, respectively. **c**, The LightRoseTTA’s prediction of CASP14 domain T1049-D1 (PDB code: 6Y4F) compared with the true (experimental) structure. **d**, The LightRoseTTA’s prediction of CAMEO target 8AIP A compared with the true (experimental) structure. **e**, The LightRoseTTA’s prediction of CAMEO target 8EI5 A compared with the true (experimental) structure. *The prediction is colored red, and the true (experimental) structure is colored gray*.

#### Predicting structures of proteins with insufficient homologous sequences

To evaluate the robustness of our LightRoseTTA to proteins with insufficient homologous information, we specifically test the performance on four public datasets, including Orphan and De novo (human-designed) in (Chowdhury et al., 2022) as well as Orphan25 and Design55 in (Wang et al., 2022). In these four datasets, protein samples generally possess rather limited or even no homologous sequences. To deal with these homolog-insufficient proteins, two natural language model-based methods named RGN2 (Chowdhury et al., 2022) and trRosettaX-single (Wang et al., 2022) are previously proposed, respectively. Both of them are MSA-free, which means that no homologous information is used. For a comprehensive comparison on the four datasets, we compare our results with a representative method named RoseTTAFold (Baek et al., 2021), as well as the two natural language model-based methods RGN2 and trRosettaX-single. For the Orphan and De novo (human-designed) datasets in (Chowdhury et al., 2022), the comparison results are shown in Fig. 3a-b ^1^. Our LightRoseTTA out-performs all these comparison methods, i.e., RoseTTAFold, trRosettaX-single and RGN2, on both Orphan and De novo datasets, with both higher values of TM-score and GDT TS. Moreover, the results on the recent Orphan25 and Design55 datasets (Wang et al., 2022) are shown in Fig. 3c-d. Accordingly, our LightRoseTTA achieves the best prediction performance on Orphan25 while gets lower performance only than RoseTTAFold on Design55. The comparison results verify the robustness of our LightRoseTTA to proteins with insufficient homologous information. Moreover, to intuitively show the high structural consistency between the predicted and experimental structures, we visualize several examples of LightRoseTTA-predicted structures in Supplementary Fig. 3.

**Figure 3.**
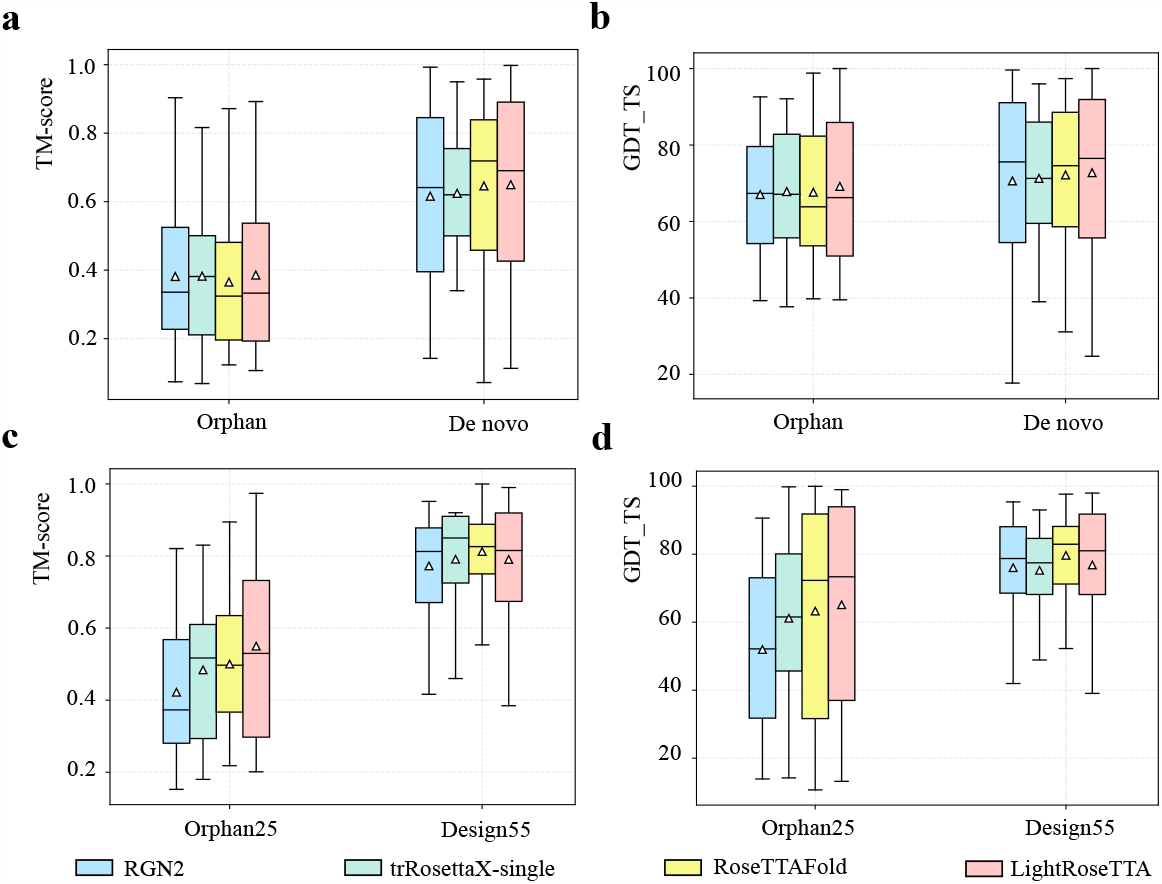
The performance comparison on the Orphan, De novo, Orphan25, and Design55 datasets. **a**, The performance (TM-score) on the Orphan and De novo datasets. For each box in the figure, the center line, bottom line, and top line represent the median, first quartile, and third quartile, respectively. The horizontal lines along the top and bottom edges represent the maximum and minimum observations. Besides, the white triangle represents the average value. For the Orphan dataset, the mean values of TM-score for RGN2, trRosettaX-single, RoseTTAFold, and LightRoseTTA are 0.381, 0.382, 0.365, and 0.385, respectively. For the De novo dataset, the values of four methods are 0.615, 0.622, 0.642, and 0.649, respectively. **b**, The performance (GDT TS) on the Orphan and De novo datasets. For the Orphan dataset, the mean value of GDT TS for RGN2, trRosettaX-single, RoseTTAFold, and LightRoseTTA are 67.04, 67.83, 67.64, and 69.15, respectively. For De novo dataset, the value of four methods are 70.63, 71.65, 72.18, and 72.82, respectively. **c**, The performance (TM-score) on the Orphan25 and Design55 datasets. For the Orphan25 dataset, the mean values of TM-score for RGN2, trRosettaX-single, RoseTTAFold, and LightRoseTTA are 0.422, 0.482, 0.491, and 0.549, respectively. For the Design55 dataset, the values of fourmethods are 0.772, 0.783, 0.812, and 0.787, respectively. **d**, The performance (GDT TS) on the Orphan25 and Design55 datasets. For the Orphan25 dataset, the mean values of GDT TS for RGN2, trRosettaX-single, RoseTTAFold, and LightRoseTTA are 51.96, 61.09, 63.25, and 65.11, respectively. For the Design55 dataset, the values of four methods are 76.08, 75.24, 79.63, and 76.57, respectively.

#### Transferring LightRoseTTA to predict antibody on Rosetta Antibody Benchmark

The antibody is a special type of protein that plays an important role in human immunity. Different from the aforementioned proteins, the structural diversity of antibodies is primarily concentrated in those complementary determining regions (CDRs). Hence, directly using the model trained on general proteins may not obtain satisfactory performance on antibody data. To solve this problem, we transfer our LightRoseTTA to predict the antibody structure by further fine-tuning the parameters based on antibody data, resulting in the model named LightRoseTTA-Ab. We test the LightRoseTTA-Ab on the Rosetta Antibody Benchmark dataset (Ruffolo et al.), and compare the performance with the results of DeepAb (Ruffolo et al.), IgFold (Ruffolo and Gray, 2022) and Ablooper (Abanades et al., 2022). For the model fine-tuning, we use the same training data with DeepAb (Ruffolo et al.).

To measure the prediction performance, one specific metric for antibody named RMSD_H3_ is adopted. RMSD_H3_ measures the RMSD value (the lower the better) of the third complementary determining region (CDR) ring of the heavy chain (CDR-H3). Specifically, the CDR-H3 is a crucial region for antibodies and presents high structural diversity, which makes structural prediction rather difficult. Fig. 4 shows the experimental results of different methods, as well as the visualized examples of predicted antibody structures. Our LightRoseTTA-Ab obtains the RMSD_H3_ value of 1.9 Å, while the results of another two state-of-the-art methods, i.e., Ablooper and DeepAb, are about 2.4 Å, and the RMSD_H3_ value of IgFold is about 3.0 Å. The lowest RMSD_H3_ demonstrates the effectiveness of our LightRoseTTA-Ab in promoting the structure prediction of antibodies. Moreover, we also visualize some prediction CDR-H3 examples of antibodies in Fig. 4b-e. As it is shown, the structural consistency of CDR-H3 can be observed between the structure predictions (red) and experiments (gray), with the RMSD_H3_ values about 1.5-1.9 Å. In addition, the RMSD_*C*α_ results of all six CDRs are shown in Supplementary Fig. 4.

**Figure 4.**
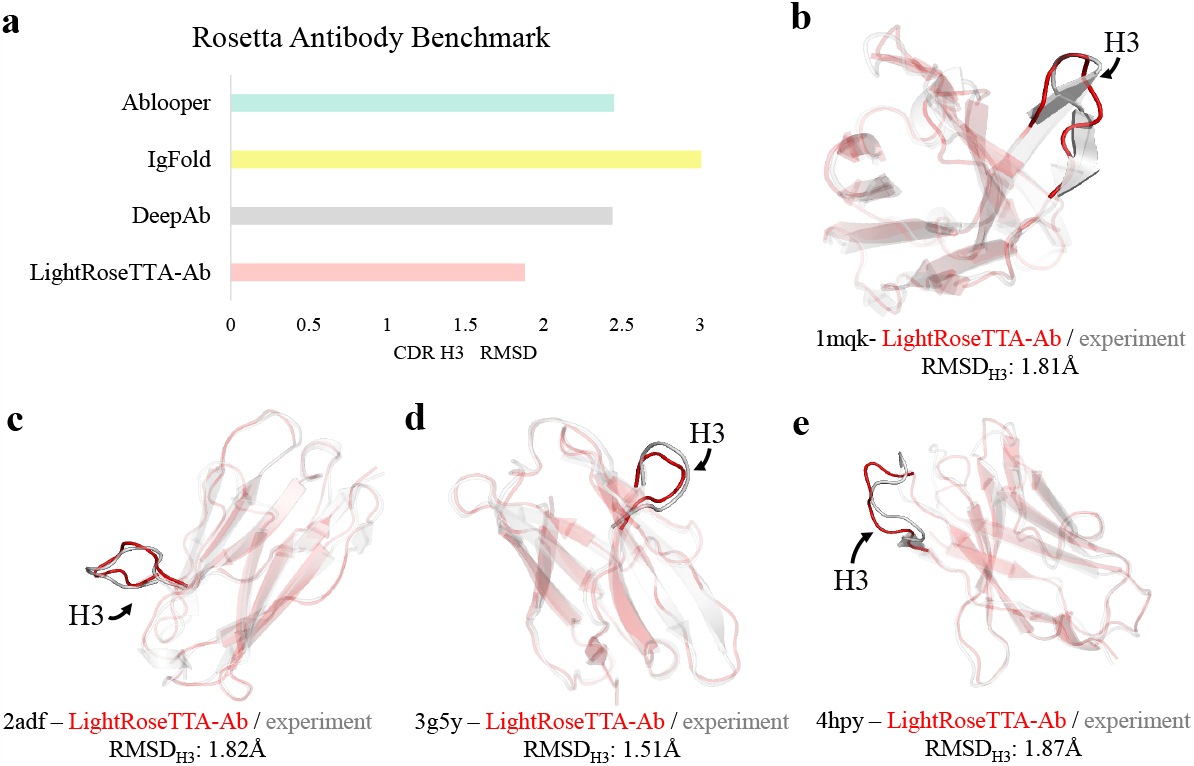
The performance comparison (RMSD-Cα of CDR-H3) on the Rosetta Antibody Benchmark and the visualization examples of the LightRoseTTA-Ab’s prediction. **a**, The performance (RMSD-Cα of CDR-H3) on the Rosetta Antibody Benchmark. **b**, The LightRoseTTA-Ab’s prediction of antibody 1mqk compared with the true (experimental) structure. **c**, The LightRoseTTA-Ab’s prediction of antibody 2adf compared with the true (experimental) structure. **d**, The LightRoseTTA-Ab’s prediction of antibody 3g5y compared with the true (experimental) structure. **e**, The LightRoseTTA-Ab’s prediction of antibody 4hpy compared with the true (experimental) structure. *The prediction is colored red, and the true (experimental) structure is colored gray*.

#### Ablation Study

To evaluate how each component contributes to LightRoseTTA’s performance, we conduct the ablation study on the CAMEO (Robin et al., 2021) and Orphan datasets (Chowdhury et al., 2022), and show the results in Fig. 5. The baseline takes a shallow co-evolution learning module followed by a shallow SE(3)-transformer. As observed in Fig. 5a-b, on the CAMEO dataset, adding each component can effectively promote the prediction accuracy with the TM-score improvements of 0.012∼0.023 if the BPE is first imposed. Contrastively, without applying the BPE constraint first, less performance improvement is obtained by each same component, as shown in Fig. 5b. The reason might be that the tremendous parameter searching space makes the model difficult to reach a suitable state, while the loss related to hard atom geometries (bond length, bond angle, etc) is difficult to decline as our experimental observation. Moreover, as shown in Fig. 5c-d, for the Orphan dataset with insufficient homologous information, the BPE also effectively promotes the structure prediction with the TM-score gain of about 0.02. Specifically, on the Orphan dataset, we pay more attention to the MSA dependency reduction (MDR) strategy to evaluate its effectiveness. According to Fig. 5c and Fig. 5d, the MDR improves the structure prediction performance of Orphan with the TM-score gain of about 0.03. This verifies the effectiveness of the MDR in improving the model’s robustness to homologous sequences. Overall, in Fig. 5, the ablation results verify the effectiveness of each component and the specific importance of MDR and BPE to boost the accuracy of protein structure prediction. The detailed description of each ablation component is described in Section 4.

**Figure 5.**
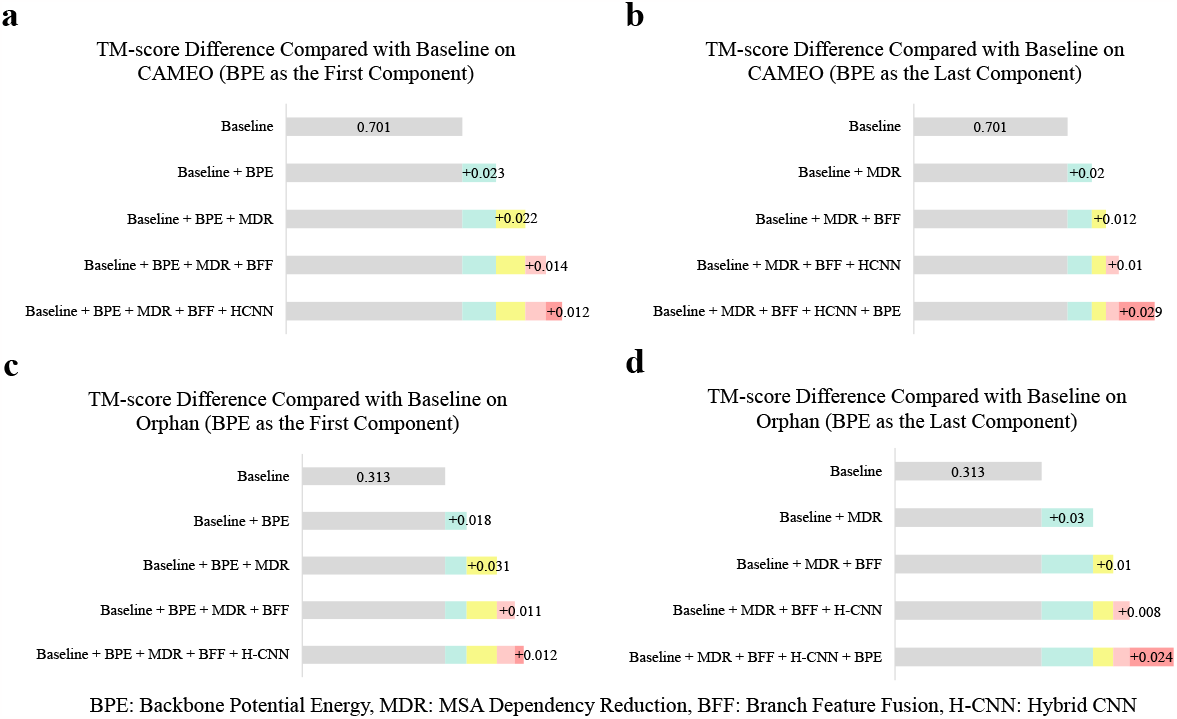
Ablation study on the CAMEO and Orphan dataset. **a**, TM-score difference between the baseline and several ablation models on the CAMEO dataset. Each ablation model is constructed by adding one or several components to the baseline. Specifically, the BPE is the first component added. **b**, TM-score difference that adds those components in a different order. Specifically, the BPE is the last-added component. **c**, TM-score difference between the baseline and several ablation models on the Orphan dataset. Specifically, the BPE is the first component added. **d**, TM-score difference that adds those components in a different order, where the BPE is the last-added component.

## 3. Discussion

In this study, the ultra-lightweight LightRoseTTA was proposed and comprehensively evaluated through both quantitative performance comparisons and visualization results. Generally, our LightRoseTTA can well handle the structure prediction for protein sequences with both sufficient and insufficient (even none) homologous sequences. Furthermore, our LightRoseTTA can be easily transferred to antibodies for structure prediction, and effectively promoted the prediction performance on CDR-H3. Besides, considering the determinative relationship of protein structure for biological function, as our LightRoseTTA’s prediction obtains high structural consistency with the true structure, we further discuss how they may contribute to the study of protein function in Section 5 of Supplementary.

More importantly, our LightRoseTTA can be deployed to the general personal computer with a single GPU for efficient training. We further discuss the comparisons of the parameter quantity, time cost for training, as well as time and memory costs for prediction. The results are shown in Fig. 6a-c, where the representative high-accuracy big model, i.e., RoseTTAFold, is compared. RoseTTAFold has the number of 130M parameters, and requires 30 days to train the framework using 8 high-speed V100 GPUs. Our LightRoseTTA has only 1.4M parameters, and costs 7 days to train the model using only one NVIDIA 3090 GPU. Hence, with the high-accurate performance guaranteed, our LightRoseTTA could largely shorten the research cycle for protein structure prediction under resource-limited environment with general devices.

**Figure 6.**
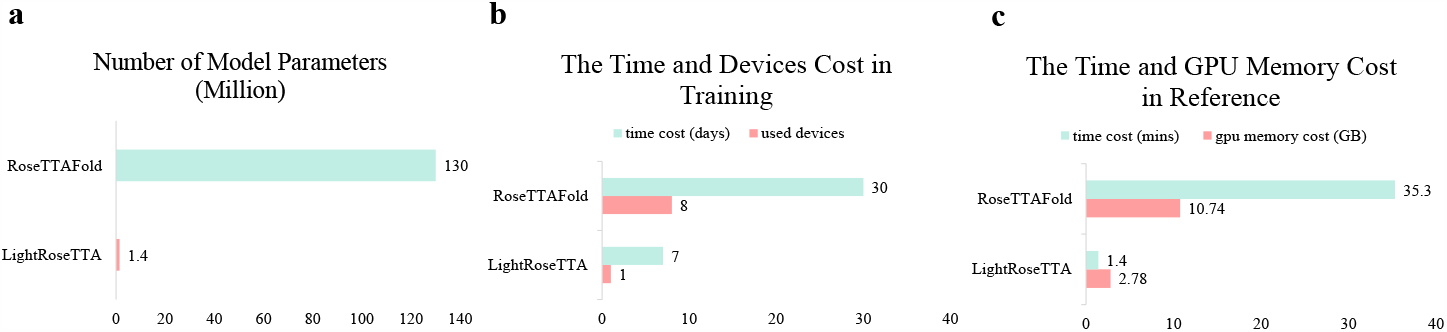
The parameter quantity, training cost, and testing cost of state-of-the-art methods and LightRoseTTA. **a**, The number of model parameters. Our LightRoseTTA has the smaller model with 1.4M parameters, while the parameter number of RoseTTAFold is 130M. **b**, The time cost and devices used for training. LightRoseTTA can be trained with a single general GPU in one week, RosettaFold costs 30 days using 8 high-speed V100 GPUs. **c**, The average time and GPU memory cost of testing one sequence on the CAMEO dataset. When predicting all-atom structures, LightRoseTTA has an average prediction time of 1.4 minutes, and RoseTTAFold needs structural refinement with pyRosetta (Chaudhury et al., 2010) and costs a running time of 35.3 minutes. For the GPU memory requirement, LightRoseTTA has the smaller value at 2.78 GB, while RoseTTAFold requires 10.74 GB.

## 4. Methods

### The model training

We train our LightRoseTTA with the protein data released in the PDB before May 1, 2018, where 40340 non-redundant protein domains are involved totally (more details of training and testing data are described in the Section 3 of Supplementary). Given the predicted structure from the 3D-Structure Generator, to guide the model optimization, the network is primarily supervised by the five-part losses, including 1) our proposed BPE (bond length *Loss*_*bb bl*_, bond angle *Loss*_*bb ba*_ and dihedral *Loss*_*bb bd*_) for the backbone generation; 2) the coordinate RMSD *Loss*_*bb RMS D*_ of the predicted backbone structure; 3) mean squared error, i.e., *Loss*_*bb pLDDT*_, of the predicted *C*_*α*_-LDDT (Jumper et al., 2021) scores of the backbone; 4) inter-residue distance and orientation losses (cross entropy), denoted as 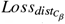, *Loss*_*ω*_, *Loss*_*θ*_, and *Loss*_*ϕ*_, respectively; 5) the prior structural losses/constraints for all-atom coordinates (bond length *Loss*_*aa bl*_, bond angle *Loss*_*aa ba*_ and dihedral *Loss*_*aa bd*_) (the details of loss functions are introduced in the Section 2 of Supplementary). Based on these losses, all model parameters can be optimized from scratch through backpropagation in an end-to-end manner. The training and testing can be run on a general PC with a single NVIDIA RTX 3090 (24GB GPU memory) card. More training details are described in the Section 1 of Supplementary.

### Input feature generation

The input features of LightRoseTTA consist of three parts, i.e., the MSA feature, the template information, and the atom-level graph representation. To extract MSA features, for each protein sequence, the HHsuite tool (Steinegger et al., 2019) is employed to retrieve homologous sequences against protein sequence database named Uniref30 (Suzek et al., 2015) (released before June 2020) and BFD (Jumper et al., 2021). Based on these homologous sequences, we construct MSA features whose each column corresponds to the position in the alignment sequence, while each row indicates each sequence in the MSA. For the template construction, we search the three-dimensional structures of homologous sequences in the structure database named PDB100 (wwPDB Consortium., 2018) (released before March 2021, as same as RoseTTAFold). In the testing process, the templates corresponding to the query sequence itself are excluded.

The atom-level graph treats atoms as nodes and between-atom bonds as edges. Each node feature involves five aspects of information: (1) the type of amino acid (20 kinds of common amino acids); (2) the type of heavy atom (C, N, O, S); (3) the index of heavy atom in amino acid; (4) the label to determine whether each atom belongs to the backbone or side chain; (5) initial 3D-coordinates of atom in the unfolded state. Among them, the first four items are represented as 20-dimensional, 4-dimensional, 14-dimensional and 2-dimensional one-hot vectors, respectively. Finally, all these parts of features are concatenated as a 43-dimensional vector.

### The residue-level branch

The residue-level branch takes the MSA features and their corresponding templates as inputs. It consists of three learning modules, including the co-evolution learning, the hybrid CNN, and the residue graph learning modules. For the co-evolution learning, first, we use the interactive update mechanism between MSAs and templates, similar to that in the RoseTTAFold (Baek et al., 2021). Different from (Baek et al., 2021), we only need to stack a shallow architecture of a few co-evolution blocks with the help of our designed BPE. After the co-evolution stage, both residue features and pairwise inter-residue relation are obtained. Then, a new hybrid CNN is designed by fusing both classic (unsymmetric) and symmetric convolutional kernels, so that the inter-residue geometries are further learnt, including the symmetric parts (inter-residue distance 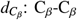 and inter-residue dihedral *ω*: C_*α*_-C_*β*_-C_*β*_-C_*α*_), and the unsymmetric parts (the interresidue dihedral *θ*: N-C_*α*_-C_*β*_-C_*β*_ and inter-residue planar angle *ϕ*: C_*α*_-C_*β*_-C_*β*_). Next, based on those residue features and their inter-residue representation, a residue-level graph is constructed by taking them as nodes and edges, respectively. Finally, a graph transformer (Shi et al., 2021) is employed for geometric representation learning, where multi-head attention coefficients are learned for node aggregation using a message passing procedure.

The architecture of our designed hybrid CNN, where two types of convolutional kernels, including both classic (unsymmetric) and symmetric ones, are used (shown in Supplementary Fig. 1a). Specifically, the symmetric kernels can be constructed through the summation of the unsymmetric kernels and their transposes. For the convolutional learning, the unsymmetric kernels are used for the inter-residue dihedral *θ* and inter-residue planar angle *ϕ*, while the symmetric kernels count for the inter-residue distance 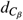 and inter-residue dihedral *ω*. By virtue of the symmetric convolution, the symmetries of those inter-residue geometries (i.e., 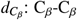 and *ω*: C_*α*_-C_*β*_-C_*β*_-C_*α*_) can be well preserved. In detail, the symmetric convolution contains 3 blocks, and each block has a 3×3 symmetric convolution kernel layer, with a BatchNorm layer and an activation exponential linear unit (ELU). Formally, the symmetric convolution can be written as:

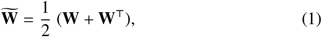

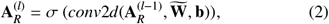

where **W** is the learnable convolution kernel with **W**^⊤^ denoting the corresponding transpose, **b** is the bias, and *σ* is the ELU activation. **A**_*R*_ denotes the matrix of inter-residule relationship, and *l* is the layer number.

### The MSA-dependency reduction strategy

To make our LightRoseTTA robust against homologous information, an MSA-dependency reduction strategy is designed for the training process. For the homologous sequences generated through the HHsuite tool (Steinegger et al., 2019), the strategy takes two MSA selection manners according to the probability of binomial distribution, where one is the top-k selection, and the other is a two-stage sampling. For instance, 30% of the training data selects the two-stage sampling result of homologous sequences as MSA input if a 0.3-probability binomial distribution is adopted. Concretely, for the top-k selection, we select 100 sequences at top in the result searched from the UniRef30 (Suzek et al., 2015) and BFD(Jumper et al., 2021) databases. For the two-stage sampling, we first generate the number of the chosen homologous sequences, then randomly sample homologous sequences. Specifically, we first set the number of chosen homologous sequences as k = Uniform[1, min(*n*_*H*_, 100)] where *n*_*H*_ is the number of all searched homologous sequences from the used homologous sequences databases. Then, we choose random k homologous sequences as the MSA input. This sampling process will increase those cases of using a limited number of homologous sequences as input during the training process. Especially when k=1, only the protein sequence itself is used as the MSA input.

### The atom-level branch

Considering that different sidechains may pose various influence to backbone conformation, we additionally construct the atom-level branch to make full use of the sidechain information. In this branch, atom-level graphs are first constructed for the common 20 kinds of amino acids by regarding atoms as nodes and bonds as edges. According to the amino acid sequence, we assemble these graphs through the peptide bond “-C-N-” on the basis of the principle of dehydration and condensation. Then, a three-layer GNN block (Morris et al., 2019) is used to learn the representation of atoms so that the sidechain information can be propagated to the backbone atoms. Each GNN layer is constructed as *X*^(*l*+1)^ = *GNN*(*A, X*^(*l*)^), where *X*^(*l*)^ represents the feature matrix at the *l*-th layer, and *A* is the adjacent matrix. Among these GNN layers, shortcut connections are added to improve the representation ability.

### The two-branch fusion

We design a variational learning process to learn weighting factors to fuse the residue-level and atom-level branches (the details are shown in Supplementary Fig. 1b). Hereby, the residue features can be refined by mining local atom-level structures that affect inter-residue bonding, e.g. torsion and folding. Here, we use 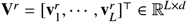 (*L* is the number of residues) to denote the residue feature matrix, and 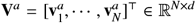 (*N* is the number of atoms) to denote the atom feature matrix. In the fusion process, the atom-residue similarity matrix **S** ∈ ℝ^*L*×*N*^ is first calculated, where the element in the *i*-th row and *j*-th column is formulated as 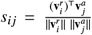. *s*_*i j*_ represents the relationship between the *i*-th residue and *j*-th atom in the protein.

Considering the complicated influence of atoms on the backbone conformation, we resort to probabilistic reasoning to select salient atoms that may influence structural bonding, so as to refine residue features. Given the atom-residue relationship **S**, we derive a set of random variables, denoted as 𝒵, by learning the posterior probability *p*(𝒵|**V**_*r*_, **V**_*a*_, **S**). Each element *z*_*i j*_ ∈ 𝒵 follows the Bernoulli distribution, denoted as *z*_*i j*_ ∼ *B*(*p*_*i j*_), where *p*_*i j*_ represents the probability that the *j*-th atom may influence the structural bonding of the *i*-th residue. However, the posterior probability *p*(𝒵|**V**_*r*_, **V**_*a*_, **S**) is usually intractable. For this problem, inspired by the reparameterization trick in (Kingma and Welling, 2013), we develop the Bernoulli reparameterization to derive the posterior probability *p*(𝒵|**V**_*r*_, **V**_*a*_, **S**). Based on 𝒵, salient atom features are selected for each residue. These selected atom features are concatenated with the corresponding residue feature, and then fed into the 3D-Structure generator for prediction.

### The 3D-Structure Generator

The 3D-Structure generator transforms the residue features to all-atom coordinates through a two-stage inference. In the first stage, the multi-layer perceptron is used to project residue features into the rough 3D coordinates, i.e., the coordinates of backbone N, *C*_*α*_ and C atoms for each residue. Then, a shallow SE(3)-transformer is employed to further refine the backbone atoms. In the second stage, the sidechain atoms are first bonded to the backbone. Then, an all-atom structural optimization mechanism, motivated by the molecular dynamics, is proposed to guide the learning of all-atom structure. Concretely, the proposed all-atom optimization mechanism consists of two parts, i.e., the prior structural constraints and the Langevin integrator (Matthews, 2015). The prior structural constraints (described as Supplementary Eqn. (3)) regularize the bond length, bond angle and dihedral of the prediction, while the Langevin integrator effectively prevents the structural distortion of the all-atom structure.

## Acknowledgements

This work was supported by the National Natural Science Foundation of China (Grant No. 62072244)

## Footnotes

1 For samples in the four datasets, homologous sequences, although quite limited or none, are still searched from Uniref30 (Suzek et al., 2015) released as of June 2020 and BFD (Jumper et al., 2021). Therefore, the reported results of RoseTTAFold in Fig. 3a-d are better than those in (Chowdhury et al., 2022; Wang et al., 2022) with no homologous sequences used.

